# Functional insight and cell-specific expression of the adipokinetic hormone/corazonin-related peptide in the human disease vector mosquito, *Aedes aegypti*

**DOI:** 10.1101/2022.07.01.498442

**Authors:** Salwa Afifi, Azizia Wahedi, Jean-Paul Paluzzi

## Abstract

The adipokinetic hormone/corazonin-related peptide (ACP) is an insect neuropeptide structurally intermediate between corazonin (CRZ) and adipokinetic hormone (AKH). Unlike the AKH and CRZ signaling systems that are widely known for their roles in the mobilization of energy substrates and stress responses, respectively, the main role of ACP and its receptor (ACPR) remains unclear in most arthropods. The current study aimed to localize the distribution of ACP in the nervous system and provide insight into its physiological roles in the disease vector mosquito, *Aedes aegypti*. Immunohistochemical analysis and fluorescence *in situ* hybridization localized the ACP peptide and transcript within a number of cells in the central nervous system, including two pairs of laterally positioned neurons in the protocerebrum of the brain and a few ventrally localized neurons within the pro- and mesothoracic regions of the fused thoracic ganglia. Further, extensive ACP-immunoreactive axonal projections with prominent blebs and varicosities were observed traversing the abdominal ganglia. Given the prominent enrichment of *ACPR* expression within the abdominal ganglia of adult *A. aegypti* mosquitoes as determined previously, the current results indicate that ACP may function as a neurotransmitter and/or neuromodulator facilitating communication between the brain and posterior regions of the nervous system. In an effort to elucidate a functional role for ACP signaling, biochemical measurement of energy nutrients in female mosquitoes revealed a reduction in abdominal glycogen stores in response to ACP that matched the actions of AKH, but interestingly, a corresponding hypertrehalosaemic effect was only found in response to AKH since ACP did not influence circulating carbohydrate levels. Comparatively, both ACP and AKH led to a significant increase in haemolymph carbohydrate levels in male mosquitoes while both peptides had no influence on their glycogen stores. Neither ACP nor AKH influenced circulating or stored lipid levels in both male and female mosquitoes. Collectively, these results reveal ACP signaling in mosquitoes may have complex sex-specific actions, and future research should aim to expand knowledge on the role of this understudied neuropeptide.

## Introduction

Neuropeptides play a crucial role in regulating numerous physiological and behavioral processes such as feeding, development, water balance, and reproduction in multicellular animals, including arthropods (Barón et al., 2010; Nässel and Winther, 2010). Three neuropeptide signaling systems present in invertebrates show homology to the mammalian gonadotropin-releasing hormone (GnRH) system (Gäde et al., 2011; Hansen et al., 2010; Li et al., 2016; Roch et al., 2011). These three neuropeptide systems include adipokinetic hormone (AKH), corazonin (CRZ) and the adipokinetic hormone/corazonin-related peptide (ACP) family.

AKH is one of the first insect neuropeptides to have been purified and isolated and is synthesized exclusively by neurosecretory cells (NSC) of the corpora cardiaca (CC), a small neurohaemal organ that is associated closely with the brain in insects (Diederen et al., 1987; Diederen et al., 2002; Stone et al., 1976). AKH was termed “adipokinetic” due to its primary function in regulating energy homeostasis in adult insects by mobilizing lipids and/or carbohydrates from the fat body (an organ equivalent to the vertebrate liver and adipose tissue) during high physical activities, including locomotion and flight (Gäde et al., 1997; Liu et al., 2009; Mercier et al., 2007). Additional roles of the AKHs include the regulation of oxidative stress, life span extension, heart-beat rate stimulation, and protein synthesis inhibition (Gäde and Marco, 2006; Zandawala et al., 2018).

Another closely related insect neuropeptide, CRZ (from corazón, which is Spanish for “heart”), also shares some structural homology with AKH but differs notably in its functions. CRZ is produced chiefly in the neuroendocrine cells of the pars lateralis of the protocerebrum and is released via the CC (Predel et al., 2007). Immunohistochemical studies in the cockroach *Periplaneta americana* (Veenstra and Davis, 1993) and fruit fly *Drosophila melanogaster* (Choi et al., 2005) showed that CRZ localized to several dorsolateral neurons in the pars lateralis and also in each abdominal ganglia of the ventral nerve cord. CRZ was first discovered in the cockroach, *P. americana*, and was named because of its cardio-excitatory activity in many insects, including the kissing bug *Rhodnius prolixus* (Patel et al., 2014; Veenstra, 1989). However, CRZ lacks cardio-excitatory activity in adult *Anopheles gambiae* mosquitoes (Hillyer et al., 2012). Whether CRZ has a conserved function across insects remains unclear, although it has also been linked to the control of melanization, ecdysis as well as responses to metabolic and osmotic stress (Kim et al., 2004; Kubrak et al., 2016; Tawfik et al., 1999; Veenstra, 1989; Zandawala et al., 2021).

A third structurally-related signaling system was discovered in the *A. gambiae* mosquito named adipokinetic hormone/corazonin-related peptide (ACP), which is found in diverse insects and is evolutionarily related to AKH and CRZ (Hansen et al., 2010). Although ACP and its receptor (ACPR) are structurally intermediate between AKH and CRZ and their receptors, their functional role in insects is not well established. Nonetheless, studies revealed that AKH, CRZ, and ACP signaling systems function independently as their receptors show highly-selective specificity for their respective ligands (Hansen et al., 2010; Zandawala et al., 2015; Hamoudi et al., 2016; Oryan et al., 2018; Wahedi and Paluzzi, 2018). Moreover, studies utilizing comprehensive *in silico* analyses proposed that the AKH hormonal system was duplicated prior to the emergence of the phylum Arthropoda, resulting in the ACP and AKH signaling systems (Hansen et al., 2010; Hauser and Grimmelikhuijzen, 2014; Marchal et al., 2018; Zhou et al., 2018).

Studies in multiple insects, including the kissing bug *R. prolixus* and female mosquito *A. gambiae*, have shown that ACP does not regulate the mobilization of energy substrates and thus does not replicate the actions of AKH (Kaufmann and Brown, 2008; Patel et al., 2014). Furthermore, ACP does not influence the heart-beat rate, indicating that the physiological actions of ACP do not overlap with a prominent function of CRZ (Hillyer et al., 2012; Patel et al., 2014). Interestingly, a precise function of ACP was found recently where it regulates haemolymph levels of carbohydrates and lipids in the male cricket, *Gryllus bimaculatus* (Zhou et al., 2018). This evidence indicated that ACP in crickets might indeed share a functional role with AKH in the regulation of energy homeostasis (Zhou et al., 2018). Moreover, an elegant study in the locust *Locusta migratoria* revealed that ACP regulates muscle lipid utilization during a long-term migratory flight (Hou et al., 2021). Contrary to the utilization of lipids as a primary energy substrate in locusts, the primary source of energy used by mosquitoes is carbohydrates in the form of trehalose, although prolonged stress will also result in lipid breakdown (Briegel et al., 2001).

Transcripts encoding *ACP* and its receptor (*ACPR*) were found to be enriched in the nervous system in *R. prolixus* and *Tribolium castaneum* (Hansen et al., 2010; Zandawala et al., 2015). ACP was immunolocalized to numerous neurons in each hemisphere of the brain, with projections throughout the central nervous system (CNS) in *T. castaneum* and *R. prolixus*, but notably, no projections were found exiting the CNS (Hansen et al., 2010; Patel et al., 2014). A recent study in Orthoptera revealed that the ACP immunolocalization is surprisingly different from other insect species since ACP is produced by neuroendocrine cells in both the pars intercerebralis and pars lateralis (e.g. *L. migratoria*), while in other insect species (e.g. *P. americana*), ACP is restricted to neurons in the pars lateralis alone (Veenstra, 2021). In adult *A. aegypti, ACP* and *ACPR* transcripts are expressed in the CNS with *ACP* transcript significantly enriched in the female brain and thoracic ganglia, while the *ACPR* transcript was significantly enriched in the abdominal ganglia of both sexes (Wahedi and Paluzzi, 2018). Furthermore, the *A. aegypti ACPR* transcript was expressed in other tissues outside the nervous system, such as the carcass, which includes the fat body of male adult mosquitoes, suggesting that the ACP signaling system might also have a role in peripheral tissues (Wahedi and Paluzzi, 2018).

Consequently, the current study aimed to advance our understanding of the localization and specific physiological role of the ACP/ACPR signaling system in the adult disease vector mosquito, *A. aegypti*. This included mapping the distribution of the ACP peptide and transcript throughout the CNS and examining the *ACPR* transcript expression pattern within the ventral nerve cord comparing abundance in pre-terminal abdominal ganglia and the terminal ganglion, which would provide insight into potential functional roles. Lastly, energy substrates were quantified following injection with *A. aegypti* ACP (*Aedae-*ACP) to determine its effect (if any) on the mobilization of trehalose and lipid and their depletion from stores in both male and female adult *A. aegypti*.

## Materials and Methods

### Experimental animals and rearing conditions

*Aedes aegypti* mosquito rearing was carried out as described previously (Rocco et al., 2017; Wahedi and Paluzzi, 2018). Briefly, adults of *A. aegypti* (Liverpool strain) were obtained from an established laboratory colony in the Department of Biology, York University (Toronto, ON, Canada). Larvae of *A. aegypti* were hatched from semi-desiccated eggs oviposited onto Whatman filter papers in a plastic container filled with double-distilled water and were fed daily with several drops of a larval feed solution comprised of 2% (w/v) beef liver powder and 2% (w/v) brewer’s yeast. Pupae were transferred to beakers containing distilled water. Larvae, pupae, and adult mosquitoes were reared in an incubator (26°C, 12:12 hour light: dark cycle). Adult male and female mosquitoes were supplied with a 10% sucrose solution through a cotton ball wick fitted in a microcentrifuge tube. Colony maintenance included adult females that were blood-fed using an artificial membrane feeding system every two days with sheep’s blood in Alsever’s solution (Cedarlane Laboratories Ltd., Burlington, ON, Canada). All experiments were carried out using one to four-day-old female and male adult mosquitoes (fed *ad libitum* with 10% sucrose) that had been isolated and transferred into mesh-covered glass jars in early pupal stage.

### Immunohistochemistry

Male and female one and four-day-old adult *A. aegypti* were briefly anesthetized with CO_2_, and nervous system tissues (i.e., brain, thoracic ganglia, and abdominal ganglia) were dissected at room temperature (RT) in 1x nuclease-free Dulbecco’s phosphate-buffered saline (1x DPBS) and then transferred immediately to 4% paraformaldehyde for fixation overnight at 4°C. The nervous tissues were then washed three times, 15 minutes each with 1x DPBS, and the tissues were then incubated at RT for one hour with 4% Triton X-100, 2% bovine serum albumin (BSA), and 10% normal sheep serum (NSS) prepared in 1x DPBS. Following this, tissues were washed several times with 1x DPBS, each wash lasting 15 minutes to remove all traces of permeabilization solution. The nervous tissues were incubated with an anti-ACP mouse polyclonal ACP antiserum (diluted 1:1000), which was a kind gift from Prof. Jan Veenstra (Patel et al., 2014; Veenstra, 2021), prepared in 1x DPBS with 0.4% Triton X-100, 2% BSA(w/v) and 2% NSS(v/v) for 4 days with gentle agitation on a flatbed rocker at 4 °C. As a negative control, the control tissues were incubated under the same conditions and in the same solution [0.4% Triton X-100 containing 2% BSA (w/v) and 2% NSS (v/v) in 1x DPBS] but without adding the ACP primary antiserum. After the four-day ACP primary antiserum incubation, tissues underwent three 15-minute washes with 1x DPBS. The nervous tissues were then incubated overnight at 4°C with Alexa Fluor 594-Goat anti-mouse IgG (H+L) secondary antibody (1:200 dilution; Molecular Probes, Life Technologies, Eugene, OR) made up in 1x DPBS with 10% NSS(v/v). Following overnight incubation at 4°C with gentle agitation, tissues were rinsed several times with 1x DPBS at RT and mounted on glass coverslips with mounting media [1x DPBS with 50% glycerol containing 4 μg/mL 4′,6-Diamidino-2-phenylindole dihydrochloride (DAPI)] and imaged on a Nikon Eclipse T*i* fluorescence microscope (Neville, NY).

### Preparation of Digoxigenin-labeled RNA probes

The distribution of cells expressing the *A. aegypti ACP* mRNA within the CNS was determined using fluorescent *in situ* hybridization (FISH) following a similar protocol as previously described (Rocco and Paluzzi, 2020; Sajadi et al., 2020). To synthesize the sense and antisense probes, a 333bp fragment of the *A. aegypti ACP* transcript including the complete open reading frame was amplified via PCR (see **Table S1** for primer details) using whole mixed-sex adult cDNA as a template. Primer sequences for *AedaeACP* (Genbank Accession Number: FN391984) are based on a previously published sequence (Kaufmann et al., 2009). The amplicon was ligated to pGEM-T vector (Promega, Madison, WI). The T7 promoter sequence (5’-AATTGTAATACGACTCACTATAGGGCG-3’) at the 5’ end of the sense strand and the 5’ end of the antisense strand was added to the PCR products via directional screening and subsequent amplification of *ACP* cDNA from the pGEM-T vector using a combination of a T7 promoter sequence primer and either a gene-specific forward primer for antisense probe template or a gene-specific reverse primer for sense probe template.

Subsequently, digoxigenin (dig) labeled antisense and sense *ACP* probes were generated by *in vitro* transcription using the T7 RNA Polymerase Mix and 10X Reaction Buffer from the HiScribe T7 High Yield RNA Synthesis Kit (New England Biolabs, Whitby, ON) and the DIG RNA Labeling Mix, 10X conc. (Roche Applied Science, Mannheim, Germany), following the manufacturer’s protocol. Once DIG-labeled RNA synthesis was complete, template DNA was removed with DNase I (New England Biolabs, Whitby, ON) and run on a non-denaturing 1% agarose gel to confirm RNA probe integrity. RNA probes were quantified by UV spectroscopy using a Take3 micro-volume plate and measured on a Synergy Multi-Mode Microplate Reader (BioTek, Winooski, VT, USA).

### *ACP* transcript localization using fluorescence *in situ* hybridization

Four-day-old adult *A. aegypti* tissues/organs were dissected in 1x nuclease-free DPBS and immediately placed in 0.2mL PCR tubes containing fixation solution (4% paraformaldehyde) and incubated at room temperature (RT) for 60 minutes on a rocker. Tissues were subsequently washed five times with 0.1% Tween-20 in DPBS (PBT) and treated with 1% H_2_O_2_ (diluted in DPBS) for 10 minutes at RT to quench endogenous peroxidase activity. Tissues were then treated with 4% Triton X-100 (Sigma Aldrich, Oakville, Ontario, Canada) prepared in PBT and incubated for one hour at RT to digest the tissues. Tissues were then washed three times with PBT to stop the digestion. A secondary fixation was performed for 20 minutes in fresh fixation solution described above and then washed three times with PBT to remove all traces of fixative. Tissues were then rinsed in a 1:1 mixture of PBT-RNA hybridization solution (50% formamide, 5x SSC, 100 μg/mL heparin, 100 μg/mL sonicated salmon sperm DNA and 0.1% Tween-20), which was subsequently removed and replaced with 100% hybridization solution and tissues held at RT. Aliquots (100 μL/tube of samples) of RNA hybridization solution were boiled at 100°C for five minutes and then cooled on ice for five minutes, giving rise to the prehybridization solution. The tissues were incubated in the prehybridization solution at 56°C for 60 minutes. During prehybridization, 200ng of antisense RNA probe for *ACP* (or sense RNA probe for controls) was added per 100μL of fresh hybridization solution, and this solution was heated to denature at 80°C for 3 minutes and then cooled on ice for 5 minutes. The prehybridization solution was then removed, and tissues were incubated in the hybridization solution containing a denatured probe for approximately 16 hours at 56°C in a thermocycler block.

The next day, tissues were washed twice with fresh hybridization solution and subsequently with 3:1, 1:1 and 1:3 (vol/vol) mixtures of hybridization solution-PBT (all pre-warmed to 56°C). The samples were then washed with PBT pre-warmed to 56°C and cooled to room temperature. To reduce non-specific staining, tissues were blocked for one hour with PBTB (1x DPBS, 0.1% Tween-20, 1% Molecular Probes block reagent; Invitrogen, Carlsbad, CA). The tissues were then incubated in a solution containing PBTB and a 1:400 dilution of a mouse anti-DIG biotin-conjugated antibody (Jackson ImmunoResearch Laboratories, West Grove, PA) at RT for 1.5 hours on a rocker and protected from light. The antibody incubation was followed by several washes in PBTB over the course of 1 hour. Tissues were then incubated in a 1:100 dilution of horseradish peroxidase-streptavidin solution (Molecular Probes, Life Technologies, Eugene, OR) in PBTB for 1 hour, and the tissues were washed with PBTB several times over an hour. Then, the tissues were washed twice with PBT and once with DPBS. Afterwards, a tyramide solution was prepared consisting of Alexa Fluor 568 tyramide dye (1:200) in amplification buffer containing 0.0015% H_2_O_2_ (Life Technologies, Eugene, OR). After the last DPBS wash was completely removed from the tissues, the tyramide solution was added to the tissues and incubated in the dark for 1 hour on a rocker at RT. The tyramide solution was then removed, and the tissues were washed with DPBS ten times over the course of an hour. Tissues were mounted on coverslips with mounting media comprised of DPBS with 50% glycerol containing 4 μg/mL 4′,6-Diamidino-2-phenylindole dihydrochloride (DAPI) and were visualized on a Nikon Eclipse T*i* fluorescence microscope (Neville, NY).

### Abdominal and terminal ganglia dissections, RNA extraction, cDNA synthesis and RT-qPCR

One-day-old adult *A. aegypti* male (n□=□50-60) and female (n□=□40-50) in each biological replicate were submerged in 1x DPBS after immobilization with brief CO_2_ exposure. In adult mosquitoes, only six abdominal ganglia appear as a result of the fusion of the 1^st^ abdominal ganglion to the posterior of the metathoracic ganglion, and the 7^th^ and 8^th^ ganglia are fused into a terminal ganglion (Brown and Cao, 2001). Thus, the abdominal ganglia (2^nd^-6^th^) and the fused terminal ganglion were dissected separately and stored in DNA/RNA protection reagent (New England Biolabs, Whitby, ON). Total RNA was isolated and purified using the Monarch® Total RNA Miniprep Kit following the manufacturer’s protocol and guidelines (New England Biolabs, Whitby, ON). Purified total RNA samples were quantified with a Take3 micro-volume plate and measured on a Synergy Multi-Mode Microplate Reader (BioTek, Winooski, VT, USA). To assess *ACPR* transcript levels, cDNA was synthesized using the iScript™ Reverse Transcription Supermix for RT-qPCR (Bio-Rad, Mississauga, ON) following manufacturer recommendations, including a ten-fold dilution of cDNA following synthesis. The *ACPR* transcript abundance was quantified on a StepOnePlus™ Real-Time PCR system (Applied Biosystems, Carlsbad, CA) using PowerUP™ SYBR® Green Master Mix (Applied Biosystems, Carlsbad, CA). The conditions of thermal cycling were as follows: (1) UDG activation at 50□°C for 2□minutes, (2) 95□°C for 2□minutes, and (3) 40 cycles of (i) 15□seconds at 95□°C and (ii) 1□minute at 60□°C. Gene-specific primers amplifying *AedaeACPR* mRNA sequence were described previously (Wahedi and Paluzzi, 2018). Expression levels were normalized to the geometric mean of Ribosomal protein 49 (rp49), 60S Ribosomal Protein S18 (rpl8), and 40S Ribosomal Protein L8 (rpS18) housekeeping genes that were determined previously as optimal endogenous controls (Paluzzi et al., 2014). The transcript levels were plotted relative to the abdominal ganglia. *AedaeACPR* expression profile was determined using 4-5 biological replicates each including triplicate technical replicates and no-template negative controls. Data were analyzed using a two-tailed t-test (p□<□0.05) GraphPad Prism 8.02 (GraphPad Software, San Diego, USA) following log transformation.

### Intrathoracic peptide injections and biochemical analyses

To assess the influence of ACP on energy substrate levels in the adult *A. aegypti*, synthetic *Aedae*-ACP and *Aedae*-AKH (see **Table S2**, GenScript, Piscataway, NJ, USA) stocks (1mM) originally dissolved in dimethyl sulfoxide (DMSO) were diluted in 1x DPBS. A physiologically-relevant dose of 10 pmol for each peptide was injected into adult mosquitoes as this amount of *Aedae*-AKH was observed previously to elicit a hypertrehalosemic effect in adult *A. gambiae* (Kaufmann and Brown, 2008). To perform injections, a Nanoject III Programmable Nanoliter Injector (Drummond Scientific Company, Broomall, PA, USA) was fitted with fine-tipped glass needles made with a micropipette puller. Female and male one to four-day-old sugar-fed adult *A. aegypti* mosquitoes were injected with *Aedae*-ACP (or *Aedae*-AKH as a positive control) into the dorsal lateral thorax near the base of the wing. For sham-injected controls, an equivalent volume of DMSO diluted identically in 1x DPBS was injected. At 90 minutes post-injection, two abdomens from females and three abdomens from males were pooled. Each mosquito was carefully opened at the segmental line between the last two abdominal segments to allow haemolymph to diffuse into the 1x DPBS. Haemolymph was collected by incubating two females or three males together for 10 min in 100 μL 1x DPBS. A 90 μL aliquot of this diluted haemolymph solution was collected for the biochemical assay. The remaining abdomens were pooled and homogenized in 200 μL of 2% w/v sodium sulfate solution (2% w/v NaSO_4_).

Carbohydrate, lipid, and glycogen levels were measured using a modified version of a previously published protocol (Kaufmann and Brown, 2008; Van Handel and Day, 1988). For the separation of sugar and lipids, 1.6 mL (v:v, 1:1) of chloroform/methanol (CHCl_3_–MeOH) was added to each centrifuge tube containing either haemolymph solution or homogenized abdomens, and tubes were centrifuged at 3000 rpm for 1 min. Next, the supernatants were transferred into fresh centrifuge tubes, while the pellets were retained for glycogen analysis. Then, 600 μL of distilled H_2_O (dH_2_O) was added to the supernatant and centrifuged at 3000 rpm for 1 min. The upper portion (water/methanol, aqueous phase) was used for the sugar analysis, while the lower fraction (chloroform, organic phase) was kept for the lipid analysis. Standards ranging from 0-400 μg for lipids in chloroform and 0-200 μg carbohydrates in water were run alongside the samples. The glycogen in the precipitate and the standards were measured by adding 2 mL of anthrone reagent per tube immediately. For the sugar in the aqueous fraction, the tubes were heated on a heating block ∼ 90-110 °C until the solvent evaporated down into ∼ 0.1-0.2 mL, and 2 mL of anthrone reagent was then added per tube. Next, all tubes were heated for 17 minutes at ∼90-110 °C and then removed from the heating block and allowed to cool. The lipid portion of the samples and standards were placed at 90-110 °C to evaporate the solvent, and later 200 μL of sulfuric acid (H_2_SO_4_) was added and heated for 10 minutes in the heating block. Subsequently, 1 mL of vanillin-phosphoric acid reaction was added to each tube, which was removed from the heating block and allowed to cool. A reddish color appeared within ∼5 minutes, and the reaction was stable for up to 30 minutes.

Absorbance values for sample aliquots (100 μL/well) from processed experimental animals and standard samples were measured in 96-well plates by a microplate reader. The optical density (OD) at λ= 625 nm was determined and converted to microgram per individual female or male mosquito based on the regression analysis equations derived from the standard curves. Biochemical assays were performed over 3-5 independent biological replicates, including at least three technical replicates per sample on each plate. Statistical analyses were completed using GraphPad Prism 8.02 (GraphPad Software, San Diego, USA) utilizing a one-way ANOVA and Tukey’s multiple comparison post-test (p < 0.05).

## Results

### ACP immunoreactivity

In the adult mosquito CNS, ACP immunoreactivity was localized in two pairs of lateral cells in the brain, and their axonal projections were revealed in the anterior protocerebrum (**Fig. 1A**). Further, two immunoreactive cells were observed on the ventral side of the prothoracic segment, and one cell was faintly stained just anterior to the junction between the prothoracic segment and mesothoracic segments of the fused thoracic ganglia. Additionally, two cells were observed in the mesothoracic segment of the fused thoracic ganglia (**Fig. 1B**). Faintly staining axonal processes were observed in the thoracic ganglia that continued into the abdominal ganglia (**Fig. S1**). ACP immunoreactive processes were detected with prominent blebs and varicosities passing through each abdominal ganglia, including the terminal ganglion (**Fig. 1C, D**). A summary of ACP immunoreactivity in the CNS of adult *A. aegypti* is presented that represents multiple repetitions for each region of the nervous system (**Fig. 1E**).

**Figure 1.**
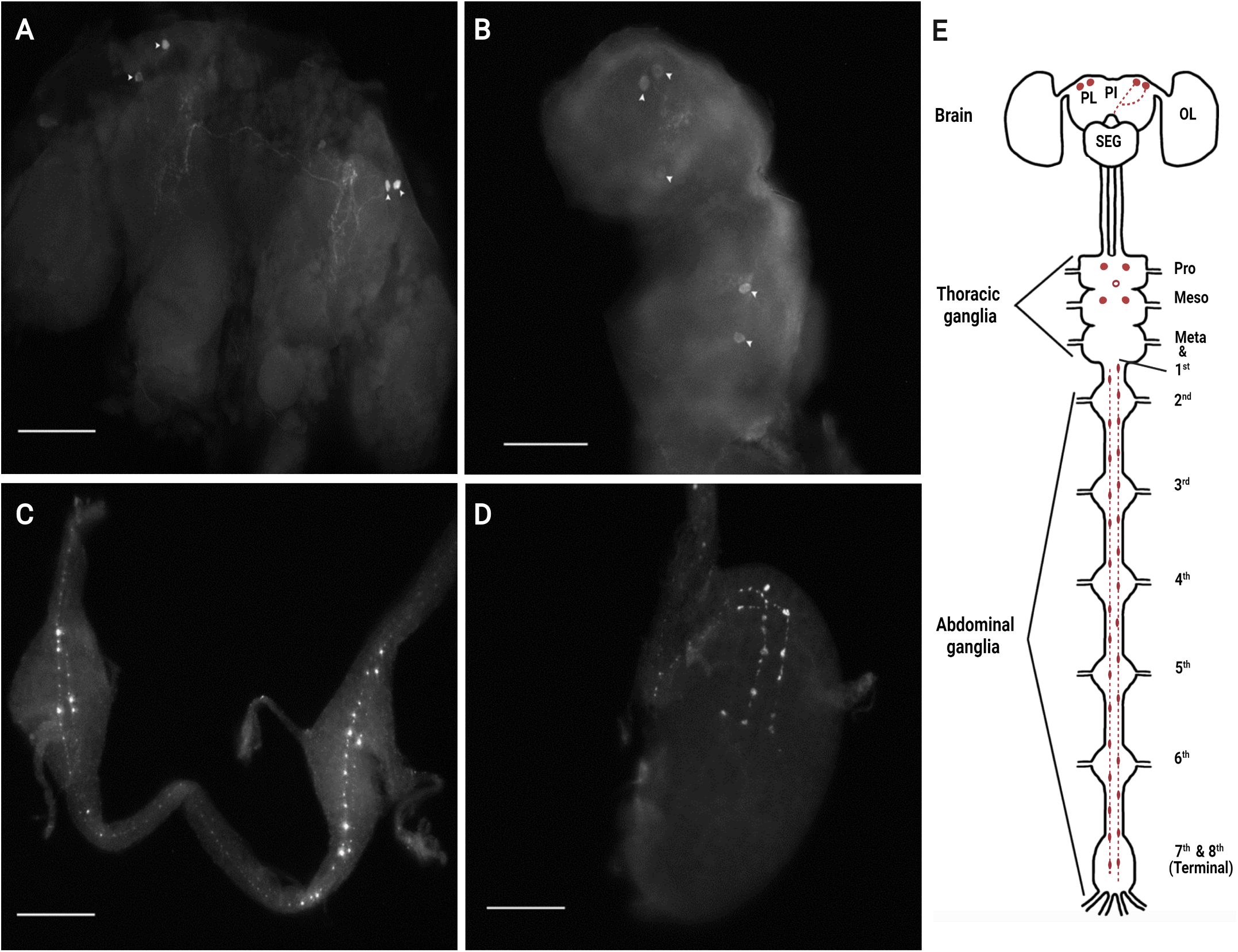
Immunolocalization of *Aedae*ACP in the nervous system of adult *A. aegypti*. ACP immunoreactivity (arrowheads) in two pairs of lateral interneurons in the brain **(A)**, a few faintly stained cells in the fused thoracic ganglia **(B)**, and extensive immunoreactive processes in all abdominal ganglia **(C)**, including the terminal abdominal ganglion **(D)**. Schematic overview illustrating the distribution of ACP immunoreactive staining in the CNS of adult *A. aegypti* **(E)**. Scale bars A and B 100 μm, C and D 50 μm. Abbreviations: (PL) pars lateralis, (PI) pars intercerebralis, (SEG) subesophageal ganglion, and (OL) optic lobe. Filled cells (•) represent a 100% detection frequency, while the unfilled cells (o) indicate inconsistent detection frequency.

### *A. aegypti ACP* transcript localization

Assessment of cell-specific expression of *A. aegypti ACP* mRNA was accomplished using fluorescence *in situ* hybridization (FISH) with antisense probes generated for the *ACP* transcript (**Fig. 2**). The CNS, including the brain, thoracic ganglia, and six abdominal ganglia of the adult ventral nerve cord, were surveyed for *AedaeACP* transcript. Similar to the ACP immunolocalization pattern, wholemounts of nervous tissues from adult male and female mosquitoes revealed fluorescence signals corresponding to *ACP* mRNA transcript in two pairs of lateral neurosecretory cells in the anterior protocerebrum and, more specifically, the supraesophageal ganglion (**Fig. 2A**). Fluorescence signals were also observed in two to three cells within the thoracic ganglia on the ventral side of the prothoracic segment just anterior to the junction between the pro-and mesothoracic segments of the ganglia (**Fig. 2B, C**). No *ACP* transcript detection was observed in the abdominal ganglia (**Fig. 2D**). No specific staining was observed in CNS tissue preparations treated with sense probes (data not shown). A schematic diagram of *ACP* transcript distribution in the CNS of adult *A. aegypti* summarizes the typical results of number and localization of *AedaeACP* transcript observed in multiple repetitions (**Fig. 2E**).

**Figure 2.**
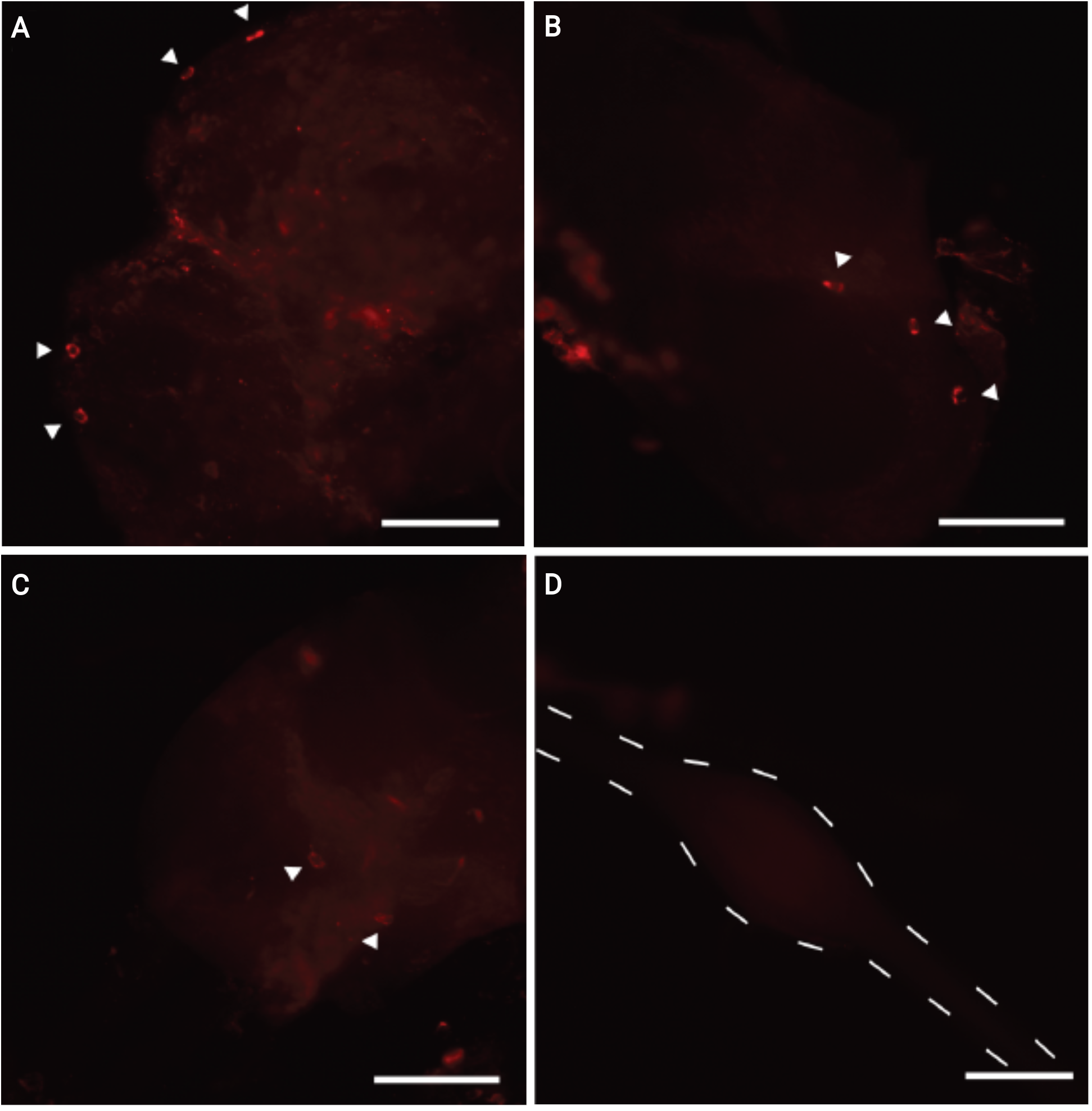
Distribution of *ACP* mRNA transcript in nervous tissue of adult *A. aegypti*. *ACP* transcript (indicated by white arrowheads) was observed in two pairs of lateral neurosecretory cells in the brain **(A)** and two to three cells in the thoracic ganglia **(B, C)**. No fluorescence was observed in the abdominal ganglia **(D)**. Scale bars A-C, 100 µm, D 50 µm.

### *ACPR* transcript abundance in the pre-terminal and terminal abdominal ganglion

RT-qPCR was utilized to determine if *ACPR* transcript was differentially expressed in the pre-terminal abdominal (2^nd^ to 6^th^) ganglia and terminal ganglion in one-day-old male and female adult *A. aegypti* mosquitoes. Notably, the expression profile of *AedaeACPR* transcript revealed no difference between the pre-terminal abdominal ganglia and the terminal ganglion in either male (**Fig. 3A**) or female (**Fig. 3B**) one-day-old adult *A. aegypti* (males, p = 0.9706 and for females, p = 0.8641).

**Figure 3.**
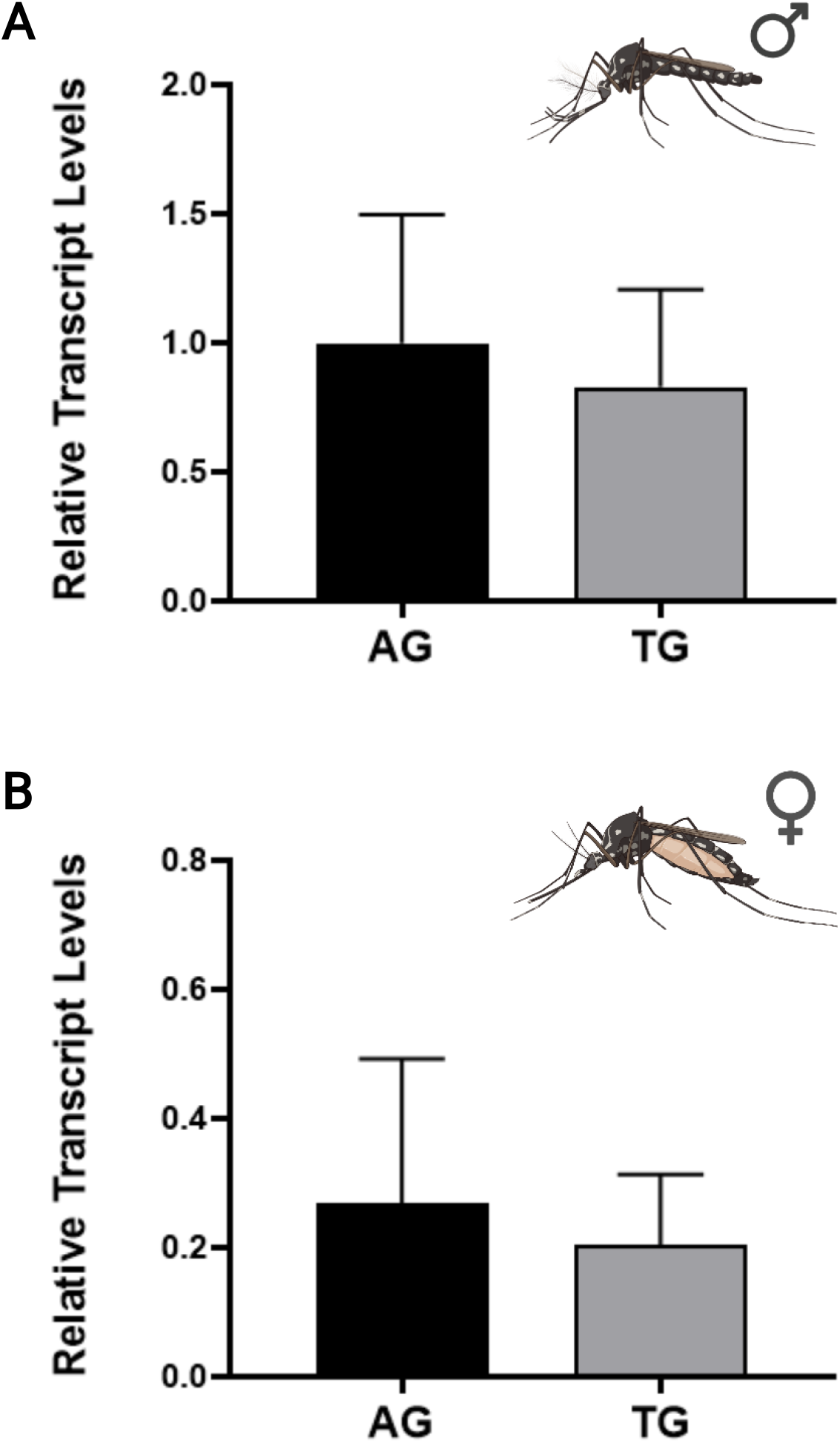
Transcript expression pattern of *ACPR* in abdominal and terminal ganglia of one-day-old adult *A. aegypti*. *ACPR* transcript abundance in **(A)** male and **(B)** female pre-terminal abdominal ganglia and terminal ganglion. Abbreviations: (AG) pre-terminal abdominal ganglia, (TG) terminal abdominal ganglion. Data represent mean□±□standard error of an average of 4-5 independent biological replicates, which was statistically analyzed using an unpaired two-tailed t-test following log transformation. The mosquito images were created using BioRender (BioRender.com).

### Metabolic effects of *Aedae*-ACP (and *Aedae*-AKH)

As a first step towards discovering physiological roles for ACP in *A. aegypti*, a potential function related to energy substrate mobilization was examined. The synthetic *Aedae-*ACP and *Aedae*-AKH were separately injected into both females (**Fig. 4**) and males (**Fig. 5**), one to four-day-old adult *A. aegypti* mosquitoes and glycogen and lipid content of abdomens and lipid and carbohydrate levels in the haemolymph were determined.

**Figure 4.**
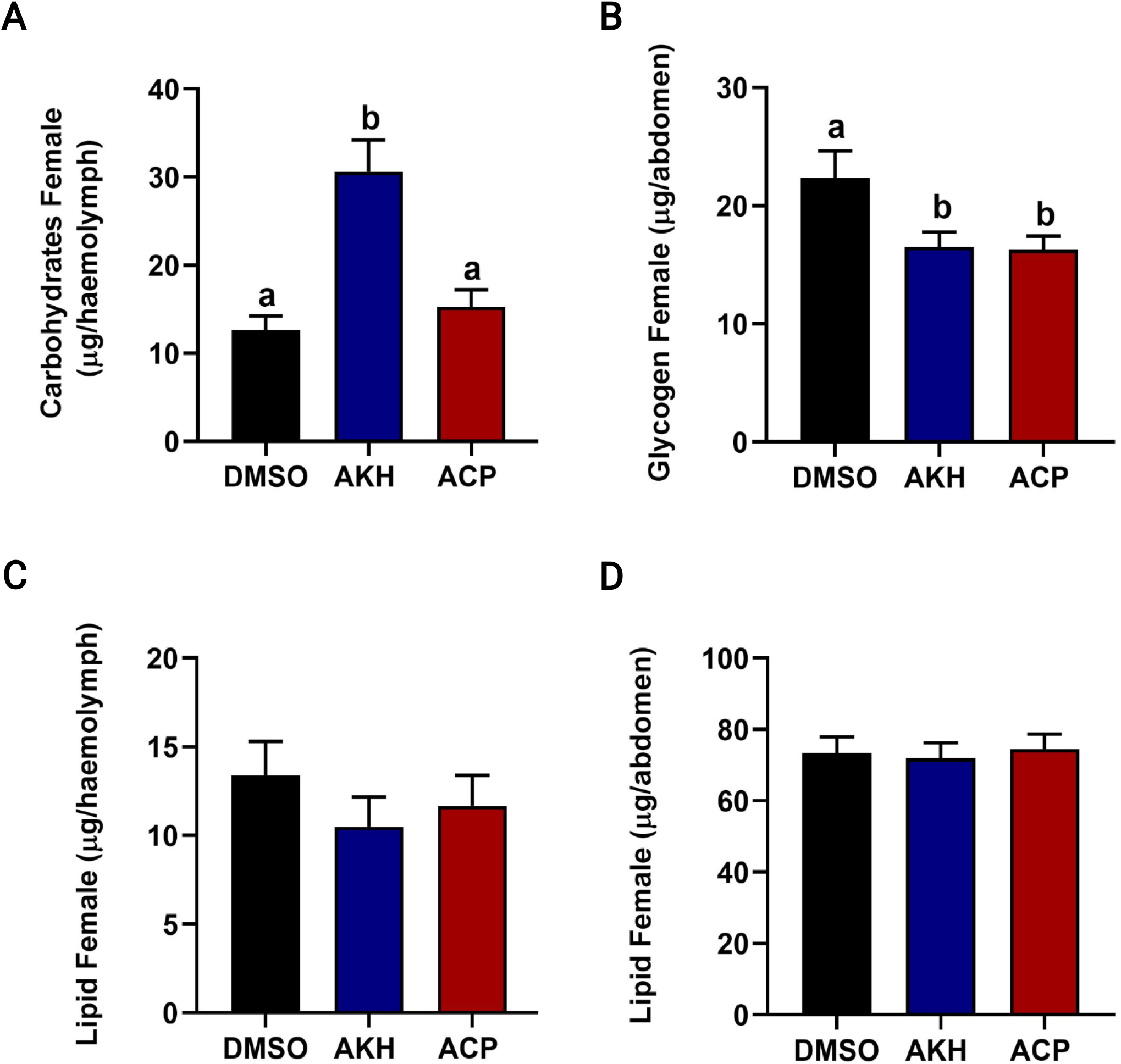
Effects of ACP and AKH on carbohydrate, glycogen, and lipid mobilization in sugar-fed female adult *A. aegypti*. **(A)** Carbohydrate in the haemolymph and **(B)** glycogen levels in the abdomen. **(C)** Lipid levels in the haemolymph and **(D)** lipid levels in the abdomen. Different letters denote bars that are significantly different from one another as determined by a one-way ANOVA and Tukey’s multiple comparison post-test (p < 0.05). Data represent the mean ± standard error of an average of 4-5 independent biological replicates.

**Figure 5.**
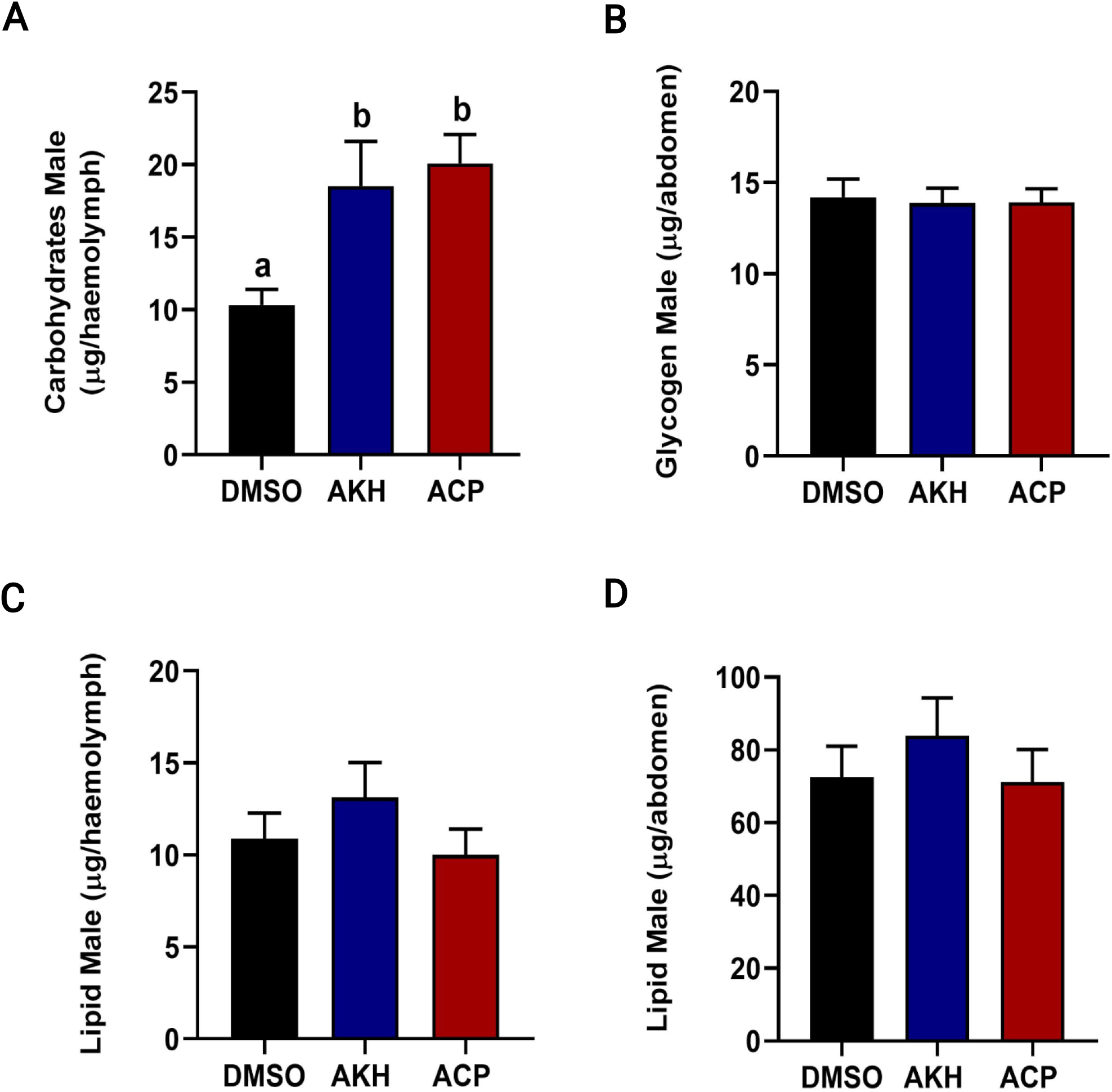
Effects of ACP and AKH on carbohydrate, glycogen, and lipid mobilization in sugar-fed male adult *A. aegypti*. **(A)** Carbohydrate in the haemolymph and **(B)** glycogen levels in the abdomen. **(C)** Lipid levels in the haemolymph and **(D)** lipid levels in the abdomen. Different letters denote bars that are significantly different from one another as analyzed by a one-way ANOVA and Tukey’s multiple comparison post-test (p < 0.05). Data represent the mean ± standard error of an average of 4-5 independent biological replicates.

At 90 min post-injection, haemolymph carbohydrate content in female mosquitoes was unchanged following *Aedae*-ACP injection but was significantly increased in females injected with *Aedae-*AKH (p < 0.0001; **Fig. 4A**). For male mosquitoes, injection with both *Aedae*-ACP and *Aedae*-AKH led to significantly elevated carbohydrate levels in the haemolymph (p = 0.0024 and p = 0.0202, respectively) (**Fig. 5A**). Glycogen level in the abdomen was significantly reduced in both *Aedae*-ACP and *Aedae-*AKH treated female mosquitoes (p = 0.0279 and p = 0.0322, respectively) (**Fig. 4B**), whereas neither peptide influenced glycogen levels in the abdomen of male mosquitoes (**Fig. 5B**). Furthermore, no significant changes in the haemolymph and abdomen lipid levels were observed after injection with *Aedae*-ACP or *Aedae-*AKH in both female (**Fig. 4C, D**) and male mosquitoes (**Fig. 5C, D**).

## Discussion

Relative to the well-studied AKH and CRZ neuropeptides, the adipokinetic hormone/corazonin-related peptide (ACP) and its receptor (ACPR) were more recently discovered and are now known to be widespread throughout many insects (Hansen et al., 2010). Although the ACP/ACPR signaling system demonstrates similarity to AKH and CRZ along with their cognate receptors, a functional relationship between these three signaling systems has not been determined, but instead, they appear to function independently (Hansen et al., 2010; Wahedi and Paluzzi, 2018; Zandawala et al., 2018). CRZ has many known functions, including cardio-stimulatory activity, melanization, ecdysis regulation, and stress physiology, while AKH is well established for its role in regulating energy mobilization in insects (Gäde et al., 1997; Kim et al., 2004; Kubrak et al., 2016; Veenstra, 1989; Zandawala et al., 2021; Ziegler et al., 1990). Studies of the ACP/ACPR signaling system in *A. aegypti* have revealed mRNA enrichment in nervous tissue, particularly in the brain and thoracic ganglia for *ACP* transcript (Kaufmann et al., 2009; Wahedi and Paluzzi, 2018) and in the abdominal ganglia for the *ACPR* transcript (Wahedi and Paluzzi, 2018).

Despite ACP and ACPR being identified in several arthropod species, unfortunately, our knowledge of their functional role in most insects remains unclear (Hansen et al., 2010; Patel et al., 2014; Zandawala et al., 2015). Therefore, this knowledge gap provides an opportunity for future investigations to better understand the specific role of this relatively understudied signaling system in insects. In the current study, the cellular localization of the ACP peptide and transcript was examined throughout the CNS of adult *A. aegypti* mosquitoes using immunohistochemistry and *in situ* hybridization. The *ACPR* transcript expression profile was determined between the pre-terminal abdominal ganglia and the terminal ganglion using RT-qPCR. Lastly, the effect of ACP on energy substrate mobilization was examined in adult female and male mosquitoes.

### Distribution pattern of ACP and ACPR in the CNS

Previous studies localized *Aedae*AKHs I and II to the brain, thoracic ganglia, and corpora cardiaca (CC) of *A. aegypti* and *A. gambiae* (Kaufmann et al., 2009; Kaufmann and Brown, 2006). *Aedae*AKH-II was later characterized as *Aedae*-ACP (Hansen et al., 2010); therefore, the immunoreactivity in cells which was herein detected in the brain and thoracic ganglia is attributed to ACP since AKH biosynthesis and storage are confined to the CC (Bogerd et al., 1995; Diederen et al., 1987; Diederen et al., 2002; Hansen et al., 2010; Kaufmann et al., 2009a; Kaufmann and Brown, 2006; Noyes et al., 1995).

ACP immunoreactivity and *ACP* transcript expression were localized in two pairs of lateral neurosecretory cells in the anterior protocerebrum of the *A. aegypti* brain. Moreover, two to three cell bodies were identified in the ventral side of the prothoracic segment; two cells were also observed in the mesothoracic segment of the fused thoracic ganglia. In addition, extensive ACP-immunoreactive axonal processes with varicosities and prominent blebs were also detected passing through each abdominal ganglion. Varicosities are involved in the cell-to-cell interactions and the formation of the presynaptic terminals that induce specific changes in their functions and structures (Giachello et al., 2012). Further, several studies found that the varicosities in the sea hare *Aplysia californica* and the snail *Helix pomatia* play a role in the neurotransmitter release modulation and growing neuron remodeling (Angers et al., 2002; Bailey and Chen, 1988; Chin et al., 2002; Cibelli et al., 1996; Ghirardi et al., 2000, 1996; Giachello et al., 2012; Hatada et al., 2000). Therefore, ACP immunoreactive varicosities and blebs associated with axonal processes observed in the abdominal ganglia provide evidence of cell-to-cell communication, suggesting this neuropeptide acts within the CNS as a neurotransmitter or neuromodulator in the mosquito. The current results are consistent (in part) with a previous study in the kissing bug, *R. prolixus*, where ACP immunoreactivity was detected in the brain, particularly in two bilaterally paired cell bodies located in the protocerebrum (Patel et al., 2014). Additionally, a recent study in the locust, *L. migratoria*, revealed that ACP peptide was detected in numerous cells in the bilateral forebrain and the pars intercerebralis (Hou et al., 2021).

Lately, a study using three orthopteran species observed that the expression of ACP is notably different from non-orthopteran insects. By examining the ACP-immunoreactivity in the adult *Schistocerca gregaria, G. bimaculatus*, and *L. migratoria*, many neuroendocrine cells were observed in the pars intercerebralis that projected to the CC, while a small number of the interneurons in the pars lateralis of the brain were detected (Veenstra, 2021). Unlike the distribution of ACP in the brain of Orthoptera, the results of the ACP immunoreactivity of the adult American cockroach *P. americana* brain (Veenstra, 2021) show consistency in the distribution of ACP within the adult brain of *A. aegypti* mosquitoes. Specifically, in the adult *P. americana*, ACP immunoreactive interneurons were expressed only in a small number of ACP interneurons in the lateral part of the brain (i.e. pars lateralis), while no ACP neuroendocrine cells in the pars intercerebralis were detected (Veenstra, 2021). Hence, based on the distribution of ACP along with the characteristics of ACP-immunoreactive staining within distinct regions of the CNS in adult *A. aegypti*, this suggests that this neuropeptide may hold a function in the coordination or communication between the anterior and posterior regions of the nervous system, suggesting a role as a neurotransmitter and/or neuromodulator.

Furthermore, previous studies prior to the discovery of the ACP system (Hansen et al., 2010) reported that AKH-like immunoreactivity was detected in 2 pairs of lateral neurosecretory cells in the brain at the anterior region of the protocerebrum in both *A. aegypti* and *A. gambiae* (Kaufmann et al., 2009; Kaufmann and Brown, 2006). However, as was suggested later, this AKH-like immunoreactivity associated with the protocerebrum is likely to represent the cross-reactivity of ACP-producing neurons since the storage and synthesis of AKH is restricted to the CC (Diederen et al., 2002; Hansen et al., 2010; Kaufmann et al., 2009; Kaufmann and Brown, 2006). Moreover, AKH-like immunoreactivity was observed within one cell in the prosegment of thoracic ganglia in ten-day-old female *A. aegypti*, while three clusters of cells in the thoracic ganglia were observed specifically within the prothoracic and mesothoracic segments of *A. gambiae* (Kaufmann et al., 2009; Kaufmann and Brown, 2006). ACP-like immunoreactivity was determined in three to four neurons in the brain within the anterior region of each hemisphere, and their axons project processes to the thoracic and abdominal ganglia in the flour beetle, *T. castaneum* (Hansen et al., 2010). ACP-like immunoreactivity wasn’t detected outside of the CNS in either *R. prolixus* or *T. castaneum*, nor was it associated with neurohaemal organs (Hansen et al., 2010; Patel et al., 2014), suggesting its physiological role is constrained within the nervous system. Similarly, the results of this study also indicate that ACP-immunoreactivity is restricted within the CNS, suggesting a functional role within the mosquito nervous system.

In a few insect species, ACP has been detected in interneurons, and no neuroendocrine cells of the pars intercerebralis projecting to the CC were observed (Hansen et al., 2010; Patel et al., 2014). Thus, the ACP immunoreactive cells and *ACP* transcript detected in the adult *A. aegypti* brain appear to be interneurons signaling to the ventral nerve cord, whereas the cells in the thoracic ganglia might signal to the periphery. There are several types of interneuronal cells, such as inter-segmental ascending and descending neurons, local amacrine neurons, and wide-ranging projection neurons (Nässel and Homberg, 2006). In addition, neuropeptides in insects are expressed by both interneurons and neuroendocrine cells. The interneurons have axons projecting within the CNS, and neuroendocrine cells have axons that leave the CNS and release their products, such as hormones, into the haemolymph (Veenstra, 2021). Moreover, neurosecretory cells are usually monopolar, which have axonal processes that are mostly projected directly to the peripheral tissues where their products are released. On the other hand, interneurons are generally involved in the regulation of nervous system-derived factors. As they could send their axonal processes forming synapses in different ganglia with a far-reaching neuron, or they might also act locally at a synapse within a single ganglion (Nässel and Homberg, 2006; Nation, 2002). In insects, several neuropeptides have been detected in the brain and the abdominal ganglia (Nässel, 2002). In contrast, limited studies exist on the presence of neurosecretory cells in thoracic ganglia; therefore, the nature of the cells that were observed in the *A. aegypti* thoracic ganglia remains unclear, and further investigations are required.

The action of ACP in the adult *A. aegypti* mosquito could be derived from either interneuron in the brain or neurosecretory cells in the thoracic ganglia (or both) and mediated via the abdominal ganglia. According to a previous study that found *ACPR* transcript enrichment in the abdominal ganglia in the adult mosquito *A. aegypti* (Wahedi and Paluzzi, 2018), this supports the notion that the abdominal ganglia might be a likely primary target of ACP action. Consequently, there is more work needed in future studies to determine the function of the ACP immunoreactive interneurons in the brain and the cells observed in the thoracic ganglia, as well as understand what triggers the activation of these neurons producing ACP. Additionally, the ACP-immunoreactive axonal processes observed might regulate or have an activity in the abdominal ganglia. This suggests that ACP might be acting as a modulator and could influence the release of other neuropeptides, including (but not limited to) crustacean cardioactive peptide, kinins, CAPA, and PK1, which have been immunolocalized to adult mosquito abdominal ganglia (Chen et al., 1994; Estévez-Lao et al., 2013; Sajadi et al., 2020). Additionally, neuropeptidomic studies have revealed members of several different peptide families are present in the abdominal ganglia, including the terminal ganglion (Predel et al., 2010). Thus, further studies are required to determine downstream targets of ACP signaling in the CNS of *A. aegypti*.

### Expression pattern of *ACPR* transcript in the abdominal ganglia

In adult mosquitoes, six distinct abdominal ganglia are present due to the fusion of the 1^st^ abdominal ganglion to the meta-thoracic ganglion, while the 7^th^ and 8^th^ ganglia are fused, forming the terminal ganglion (Brown and Cao, 2001). Expression profiles of the *A. aegypti ACPR* transcript in the 2^nd^-6^th^ abdominal ganglia and the terminal ganglion were measured to guide the functional interpretation and reveal potential roles for ACP. Examination of one-day-old *A. aegypti* male and female mosquitoes revealed no significant difference in *ACPR* transcript abundance between the abdominal ganglia (2^nd^-6^th^) and the fused terminal ganglion (7^th^ & 8^th^) in both sexes. Thus, these results corroborate observations of ACP immunoreactivity with a similar distribution in the form of axonal projections in each of the abdominal ganglia. Moreover, the widespread varicosities and blebs that were revealed indicate sites of communication (i.e. synapse) within each of the ventral nerve cord ganglia. A previous report demonstrated the enrichment of the *ACPR* transcript in the abdominal ganglia while the *ACP* transcript was most abundant in the brain, followed by the thoracic ganglia (Wahedi and Paluzzi, 2018). Together, this evidence indicates ACP may hold a neuromodulator and/or neurotransmitter function in the male and female adult *A. aegypti* mosquito. However, further studies are necessary to validate this proposed role and identify the downstream targets of ACP/ACPR signaling in the adult *A. aegypti* mosquito.

### Metabolic actions of *Aedae*-ACP

AKH and ACP are more closely related to each other compared to the CRZ system (Hansen et al., 2010). As a result of this closer structurally similarity between AKH and ACP, we sought to determine whether the metabolic function of AKH extends to ACP in adult *A. aegypti* mosquitoes. Similar to the findings in female *A. gambiae* (Kaufmann and Brown, 2008), our results confirm that injection of synthetic AKH resulted in an increase in haemolymph carbohydrate levels and a reduction in glycogen stores in female *A. aegypti* mosquitoes. Thus, in *A. aegypti*, AKH can also be designated as a hypertrehalosaemic hormone, and like other AKHs, functions comparably to the vertebrate hormone glucagon in female adult mosquitoes. In contrast, there was no change in the haemolymph carbohydrate levels in response to ACP; however, interestingly, there was a significant reduction in the glycogen stores in females. This is in contrast to previous findings in *A. gambiae*, where ACP (then referred to as AKH-II) was found to have no effect on glycogen stores (Kaufmann and Brown, 2008). To date, no study has determined whether ACP has a metabolic action in male mosquitoes. Indeed, the current data indicate that both AKH and ACP caused a significant increase in the carbohydrate level in the haemolymph, while having no influence on the glycogen stores in male mosquitoes. In addition to enrichment in the abdominal ganglia, we identified that the *A. aegypti ACPR* transcript is highly expressed (over 100-fold) in the carcass, including the fat body in adult male mosquitoes but not in female mosquitoes (Wahedi and Paluzzi, 2018). This difference in the peripheral enrichment of *ACPR* could explain the observed sex-specific metabolic actions of ACP in adult *A. aegypti*. With regards to metabolic actions in relation to lipids, our data indicate there are no changes in haemolymph or abdomen lipid levels in *Aedae*-AKH or *Aedae*-ACP injected mosquitoes, irrespective of sex. This finding is not surprising since the primary metabolic fuel in mosquitoes are carbohydrates in the form of trehalose, although prolonged movement or energy demand could also result in lipid breakdown (Briegel et al., 2001; Clements, 1992).

Several recent studies revealed that many metabolic genes and pathways contribute to sex differences (Wat et al., 2021, 2020). A study in *D. melanogaster* showed that sex-specific differences in fat storage is regulated via the AKH pathway (Wat et al., 2021). Also, a study demonstrated that the AKH receptor is involved in regulating sexual behavior and pheromone production in a sex-specific and starvation-dependent manner in *D. melanogaster* (Lebreton et al., 2016). Specifically, the AKH receptor regulates only male *D. melanogaster*, but not female, sexual behavior in a starvation-dependent manner. Males lacking AKH receptors displayed severely reduced courtship activity when starved and increased duration of mating when fed (Lebreton et al., 2016). Hence, in light of these sex-specific actions of the AKH pathway, our results suggest that the ACP signaling system may similarly have a sex-specific function in adult *A. aegypti*.

In the locust *L. migratoria*, ACP facilitates the utilization and oxidation of lipid during their long-term flight. Combining metabolomic and transcriptomic analyses, it was shown that ACP is involved in the oxidation and transport of fatty acids in flight muscles (Hou et al., 2021). In mosquitoes, the most abundant free amino acid is proline, which has been suggested as a possible flight metabolite in *A. aegypti* (Scaraffia and Wells, 2003). Thus, ACP may be involved in the conversion of energy substrates, as proline is a key intermediate in metabolism functioning as a transporter of acetyl units from the fat body to the flight muscle, where they re-enter the citric acid cycle and are oxidized to produce ATP (Scaraffia and Wells, 2003; Weeda et al., 1980). Further, several studies have implicated overlap between the biological actions of CRZ and AKH in nutritional and oxidative stress (Bednářová et al., 2015; Bharucha et al., 2008; Kubrak et al., 2016; Zandawala et al., 2021). Thus, it is possible that ACP might share a stress-related role in insects.

In conclusion, this study provides evidence for the distribution of ACP within the CNS of the adult *A. aegypti* mosquito. ACP was immunolocalized within neurons in the brain and thoracic ganglia, with axonal processes projecting into the abdominal ganglia but no evidence of processes leaving the nervous system, suggesting ACP may function as a neurotransmitter or neuromodulator. *ACPR* transcript quantification in the ventral nerve cord revealed no significant difference in expression between the pre-terminal abdominal ganglia and the terminal ganglion in both sexes. ACP led to a reduction in glycogen stores in the female adult *A. aegypti*, while an increase in the carbohydrate haemolymph level was identified in male *A. aegypti* in response to ACP injection. Notably, no changes to the abdomen and haemolymph lipid levels were observed in both sexes. Together, this study provides insight to enhance our mechanistic understanding of the broad and crucial functions of the ACP signaling system in the yellow fever disease vector mosquito *A. aegypti*, which may guide novel approaches aimed at improving existing pest control strategies to reduce the burden of these medically important disease vectors.

## Supporting information

graphical abstract

## Acknowledgments

This research was supported by an NSERC Discovery Grant to J.P.P. The authors are grateful to Dr. Jan Veenstra (University of Bordeaux) for his generous gift of the mouse polyclonal ACP antiserum used in this study.

