## Supplementary information for "Functional insight and cell-specific expression of the adipokinetic hormone/corazonin-related peptide in the human disease vector mosquito, *Aedes aegypti*"

**Table S1. Primers used to amplify *A. aegypti* ACP for FISH probe synthesis and RT-qPCR analysis.**

| Oligo Name | Sequence (5’-> 3’) | Function |
| --- | --- | --- |
| ACP-F2 | ATGTGTTCTCTAAGGCGAAATAGC | Amplification of ACP target  for FISH |
| ACP-R2 | TTACAGGTGCCCATTCGAA | Amplification of ACP target  for FISH |

**Table S2. Primary structure of peptides used for injections in the biochemical assay.**

| Peptide Name | Peptide Sequence |
| --- | --- |
| *Aedae-*ACP | **pQVTFSRDWNAa** |
| *Aedae-*AKH | **pQLTFTPSWa** |

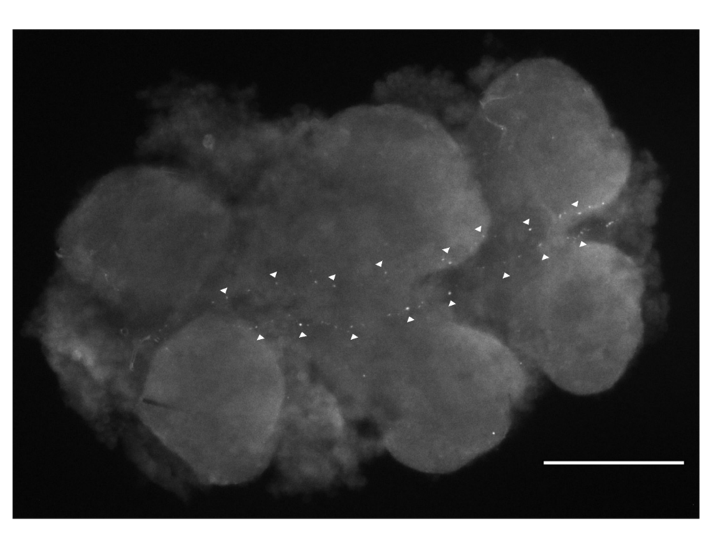

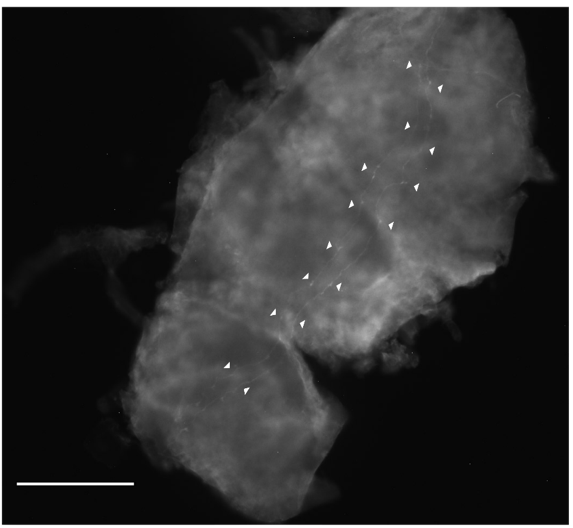

**A**

**B**

**Figure S1. ACP immunoreactive staining in axonal processes permeating the thoracic ganglia of adult *A. aegypti*.** Faintly staining ACP immunoreactive axonal processes (arrowheads) on the (A) ventral and (B) dorsal surface in the thoracic ganglia that continues into the abdominal ganglia. Scale bars 100μm.
