## Supplementary figures and images for "Functional insight and cell-specific expression of the adipokinetic hormone/corazonin-related peptide in the human disease vector mosquito, *Aedes aegypti*"

### graphical abstract

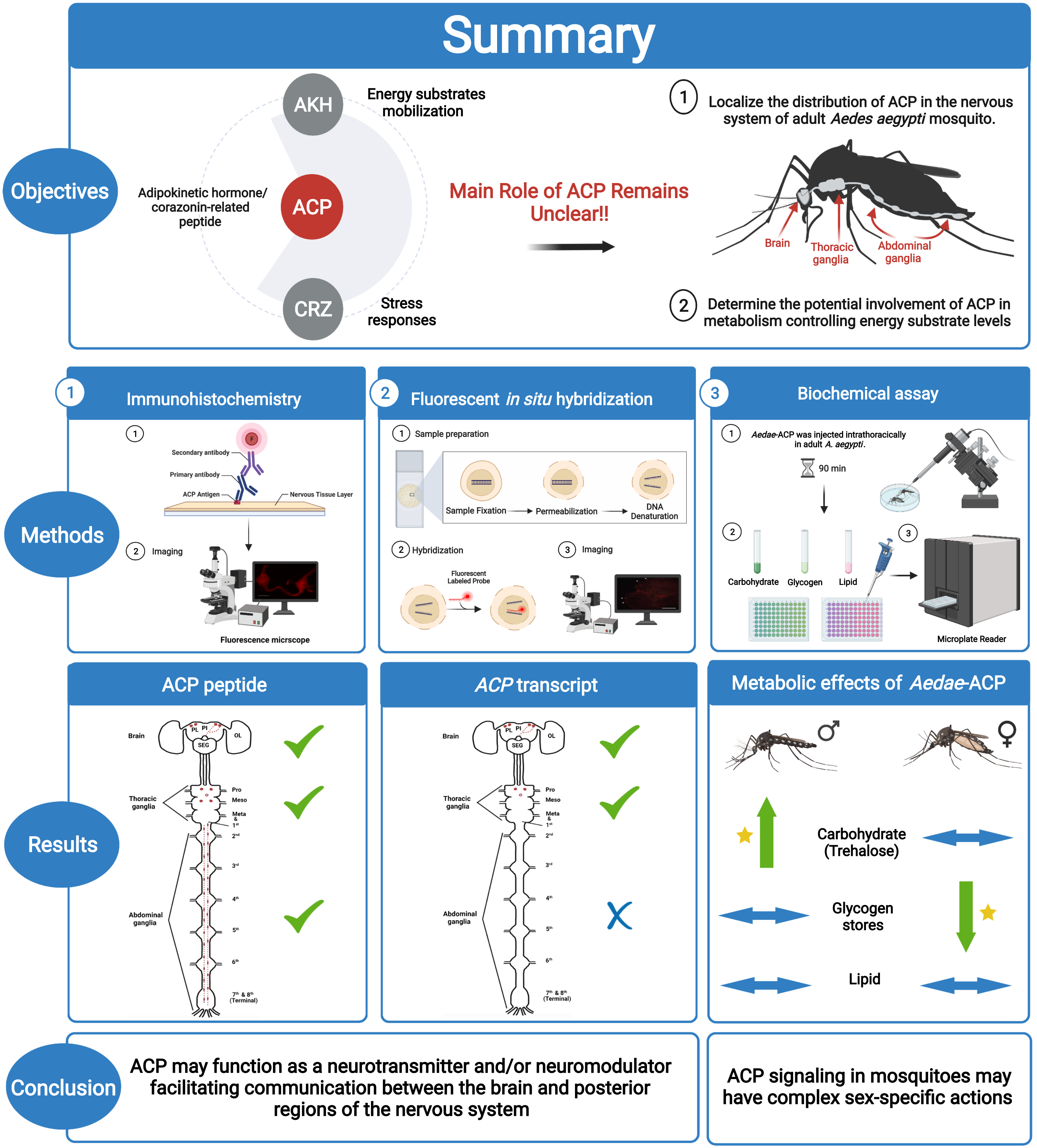
